# American Gut: an Open Platform for Citizen-Science Microbiome Research

**DOI:** 10.1101/277970

**Authors:** Daniel McDonalda, Embriette Hyde, Justine W. Debelius, James T. Morton, Antonio Gonzalez, Gail Ackermann, Alexander A. Aksenov, Bahar Behsaz, Caitriona Brennan, Yingfeng Chen, Lindsay DeRight Goldasich, Pieter C. Dorrestein, Robert R. Dunn, Ashkaan K. Fahimipour, James Gaffney, Jack A Gilbert, Grant Gogul, Jessica L. Green, Philip Hugenholtz, Greg Humphrey, Curtis Huttenhower, Matthew A. Jackson, Stefan Janssen, Dilip V. Jeste, Lingjing Jiang, Scott T. Kelley, Dan Knights, Tomasz Kosciolek, Joshua Ladau, Jeff Leach, Clarisse Marotz, Dmitry Meleshko, Alexey V. Melnik, Jessica L. Metcalf, Hosein Mohimani, Emmanuel Montassier, Jose Navas-Molina, Tanya T. Nguyen, Shyamal Peddada, Pavel Pevzner, Katherine S. Pollard, Gholamali Rahnavard, Adam Robbins-Pianka, Naseer Sangwan, Joshua Shorenstein, Larry Smarr, Se Jin Song, Timothy Spector, Austin D. Swafford, Varykina G. Thackray, Luke R. Thompson, Anupriya Tripathi, Yoshiki Vazquez-Baeza, Alison Vrbanac, Paul Wischmeyer, Elaine Wolfe, Qiyun Zhu, The American Gut Consortium, Rob Knight

## Abstract

Although much work has linked the human microbiome to specific phenotypes and lifestyle variables, data from different projects have been challenging to integrate and the extent of microbial and molecular diversity in human stool remains unknown. Using standardized protocols from the Earth Microbiome Project and sample contributions from over 10,000 citizen-scientists, together with an open research network, we compare human microbiome specimens primarily from the USA, UK, and Australia to one another and to environmental samples. Our results show an unexpected range of beta-diversity in human stool microbiomes as compared to environmental samples, demonstrate the utility of procedures for removing the effects of overgrowth during room-temperature shipping for revealing phenotype correlations, uncover new molecules and kinds of molecular communities in the human stool metabolome, and examine emergent associations among the microbiome, metabolome, and the diversity of plants that are consumed (rather than relying on reductive categorical variables such as veganism, which have little or no explanatory power). We also demonstrate the utility of the living data resource and cross-cohort comparison to confirm existing associations between the microbiome and psychiatric illness, and to reveal the extent of microbiome change within one individual during surgery, providing a paradigm for open microbiome research and education.

**Importance:** We show that a citizen-science, self-selected cohort shipping samples through the mail at room temperature recaptures many known microbiome results from clinically collected cohorts and reveals new ones. Of particular interest is integrating n=1 study data with the population data, showing that the extent of microbiome change after events such as surgery can exceed differences between distinct environmental biomes, and the effect of diverse plants in the diet which we confirm with untargeted metabolomics on hundreds of samples.

## Introduction

The human microbiome plays a fundamental role in human health and disease. While many studies link microbiome composition to phenotypes, we lack understanding of the boundaries of bacterial diversity within the human population, and the relative importance of lifestyle, health conditions, and diet, to underpin precision medicine or to educate the broader community about this key aspect of human health.

We launched the American Gut Project (AGP; http://americangut.org) in November of 2012 as a collaboration between the Earth Microbiome Project (EMP) (1) and the Human Food Project (HFP; http://humanfoodproject.com/) to discover the kinds of microbes and microbiomes “in the wild” via a self-selected citizen-scientist cohort. The EMP is tasked with characterizing the global microbial taxonomic and functional diversity, and the HFP is focused on understanding microbial diversity across human populations. As of May 2017, the AGP included microbial sequence data from 15,096 samples from 11,336 human participants, totaling over 467 million (48,599 unique) 16S rRNA V4 gene fragments (“16S”). Our project informs citizen-scientist participants about their own microbiomes by providing a standard report (fig 1A) and resources to support human microbiome research, including an online course (Gut Check: Exploring Your Microbiome; *http://www.coursera.org/learn/microbiome*). AGP deposits all de-identified data into the public domain on an ongoing basis without access restrictions (table S1). This reference database characterizes the diversity of the industrialized human gut microbiome on an unprecedented scale, reveals novel relationships with health, lifestyle, and dietary factors, and establishes the AGP resource and infrastructure as a living platform for discovery.

**Figure 1.**
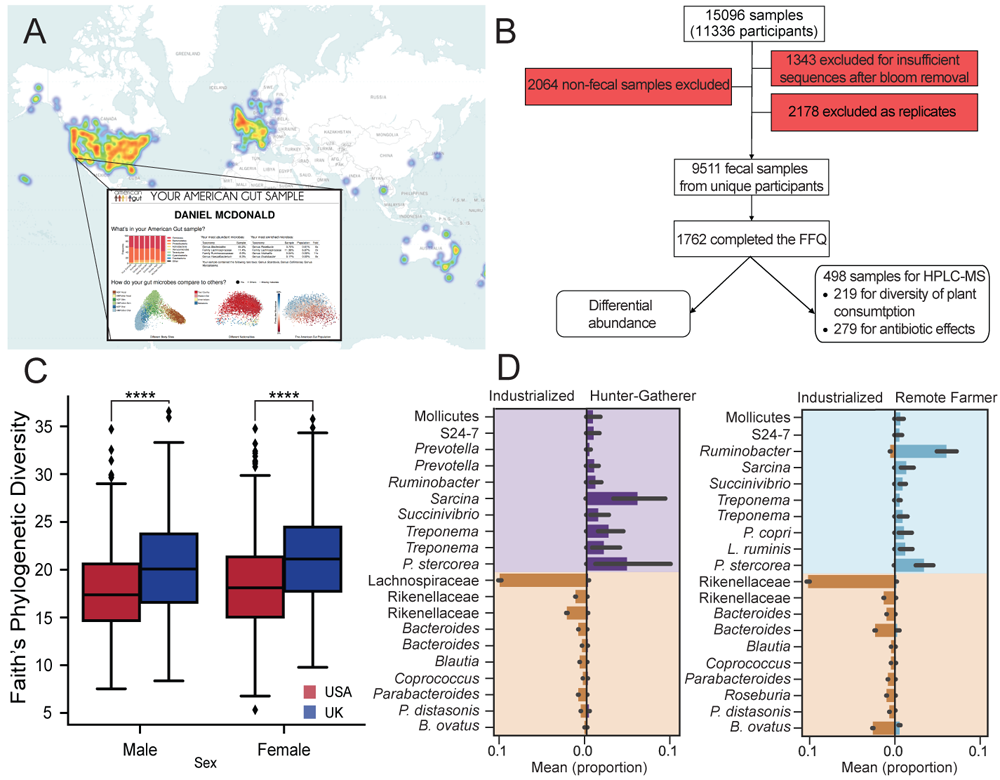
Population characteristics. **(A)** Participants across the world have sent in samples to the American Gut, although the primary geographic regions of participation are in North America and the United Kingdom; the report a participant receives is depicted. **(B)** The primary sample breakdown for subsequent analyses. Red denotes reasons samples were removed. **(C)** Between the two largest populations, the US (*n*=6,634) and the UK (*n*=2,071), we observe a significant difference in alpha diversity. **(D)** In a meta-analysis, the largely industrialized population that makes up the American Gut exhibits significant differential abundances to non-industrialized populations.

## Results

### Cohort characteristics

AGP participants primarily reside in the United States (n=7,860). However, interest in the AGP rapidly expanded beyond the US to United Kingdom (n=2,518), and Australia (*n*=321), with 42 other countries or territories also represented (fig 1A; table S1). Participants in the US inhabit urban (*n*=7,317), rural (*n*=29), and mixed (*n*=98) communities (2010 US Census data based on participant zip codes), and span greater ranges of age, race, and ethnicity than other large-scale microbiome projects (2-6). Because the AGP is crowdsourced and self-selected, and subjects generally support the cost of sample processing, the population is unrepresentative in several important respects, including having lower prevalence of smoking and obesity, higher education and income (fig S1A), and underrepresentation of Hispanic and African American communities (table S1); generalization of the results is cautioned. Targeted and population-based studies will be crucial for filling these cohort gaps (Supplemental text).

Using a survey modified from (7, 8), participants reported general health status, disease history, and lifestyle data (table S2, supplemental text). In accordance with our IRB, all survey questions were optional (median per-question response 70.9%; table S2). Additionally, 14.8% of participants completed a validated picture-based food frequency questionnaire (FFQ) (VioScreen; *http://www.viocare.com/vioscreen.html*), and responses correlated well with primary survey diet responses (table S2).

We sought to minimize errors and misclassifications well-known to occur in self-reported data (9). Survey responses relied on controlled vocabularies. For analyses, we trimmed numeric entries at extremes (e.g., weight over 200kg or below 2.5kg) and excluded obviously incorrect answers (e.g., infants drinking alcohol) and samples for which necessary data were not supplied (e.g., missing zip code data for spatial analyses); see supplement for details. We focused our primary investigative efforts on a “healthy adult” subset (*n*=3,942) of individuals aged 20-69 with BMIs ranging between 18.5–30 kg/m^2^, no self-reported history of inflammatory bowel disease, diabetes, or antibiotic use in the past year, and at least 1,250 16S sequences/sample (fig 1B, S1B).

The two largest populations in the dataset (US and UK) differed significantly in alpha-diversity, with Faith’s phylogenetic diversity (PD) higher in UK samples (*13*) (Mann Whitney *p<*1×10^−15^; fig 1C). One balance (10) (a log-ratio compositional transform) explained most of the taxonomic separation between US and UK samples (AUC=77.7% ANOVA *p*=1.01×10^−78^, *F*=386.85) (fig S1C, table S3). To understand how these two populations differed from others, we compared adult AGP samples (predominantly from industrialized regions) to samples from adults living traditional lifestyles (6, 11, 12). As previously observed (6), samples from industrial and traditional populations separated in Principal Coordinates Analysis (PCoA) space of unweighted UniFrac distances (13) (fig S1D). They show greater variation within industrial populations than within traditional populations (2) and facile separation based on microbial taxonomy (industrial vs. non-industrial agrarian: AUC=98.9%, ANOVA *p*=1.52×10^−260^, *F*=1265.8; industrial vs. hunter-gatherer: AUC=99.5%, ANOVA *p*=4.48×10^−227^, *F*=1092.35) (fig 1D, table S3).

### Removal of bacterial blooms

An important practical question is whether self-collected microbiome samples can match those from better-controlled studies. Most AGP samples are stools collected on dry swabs and shipped without preservative to minimize costs and avoid exposure to toxic preservatives. *E. coli* and a few other taxa grow in transit, so based on data from controlled storage studies as previously described (14) we removed sOTUs (sub-OTUs (15); median of 7.9% of sequences removed per sample) shown to bloom.

We further characterized the impact of these organisms through culturing, HPLC-MS analysis of cultured isolates, and shotgun metagenomics of the primary samples and storage controls (16). Culturing primary specimens stored at −80°C (US: *n=*116; UK: *n=*73; other: *n=*25) showed a strong correlation between the fraction of sequences reported as blooms in 16S sequencing and positive microbial growth following overnight incubation in aerobic conditions (fig 2A). Culture supernatants were characterized using HPLC-MS; most metabolites in these supernatants were absent from the primary specimens (fig 2B, C, method details in SI). We sequenced draft genomes of 169 isolates; of these, 65 contained the exact *E. coli* 16S sequence in the published bloom filter (14). To characterize the impact of the 16S bloom filter, we computed effect sizes over the participant covariates and technical parameters for 9,511 individual participant samples, including and excluding blooms (complete list table S2; comparisons to (17, 18) in supplementary text), and observed tight correlations for both unweighted (fig 2D, Pearson *r*=0.91, *p*=3.76×10^−57^; Spearman *r*=0.90, *p*=9.45×10^−55^) and weighted UniFrac (fig 2E, Pearson *r*=0.42, *p*=1.71×10^−6^; Spearman *r*=0.58, *p*=1.03×10^−9^). An outlier on the quantitative metric (weighted UniFrac) is present and corresponds to a variable representing the fraction of bloom reads in a sample.

**Figure 2.**
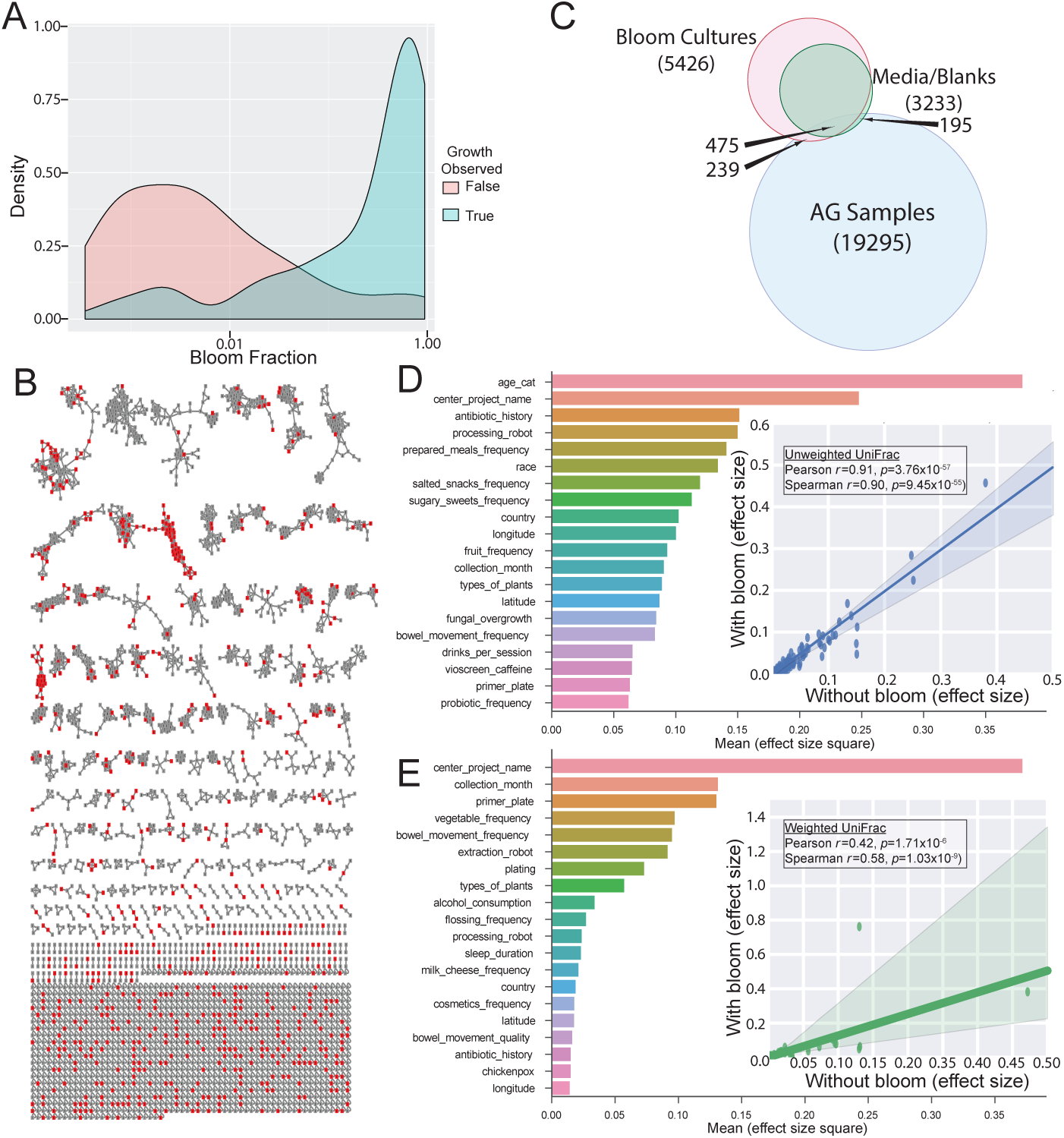
Blooms and effect sizes. **(A)** The fraction of 16S reads that recruit to bloom reads defined by Amir et al. 2017 is strongly associated with the likelihood for microbial growth under aerobic culture conditions on rich media. **(B)** Molecular network of the metabolites observed in the supernatant from cultures (*n*=217) derived from fecal samples. The nodes in red (n=239) are metabolites associated with *E. coli*. **(C)** Overlap of metabolites between AGP samples and blooms. **(D)** Unweighted UniFrac effect sizes. The inset shows the correlation of effect sizes when including or excluding the bloom 16S reads (Pearson *r*=0.91, *p*=3.76×10^−57^). **(E)** Weighted UniFrac effect sizes. The inset shows the correlation of the effect sizes when including or excluding bloom 16S reads (Pearson *r*=0.42, *p*=1.71×10^−6^); the outlier is the 16S bloom fraction of the sample.

### Novel taxa and microbiome configurations

To understand human microbiome diversity, we placed AGP samples in the context of the EMP (1). Building on earlier work revealing a striking difference between host-associated and environmental microbiomes (19), we found that the diversity of microbiomes associated with the human gut (just one vertebrate) occupies a vast extent of the microbiome diversity of the planet (fig 3A).

**Figure 3.**
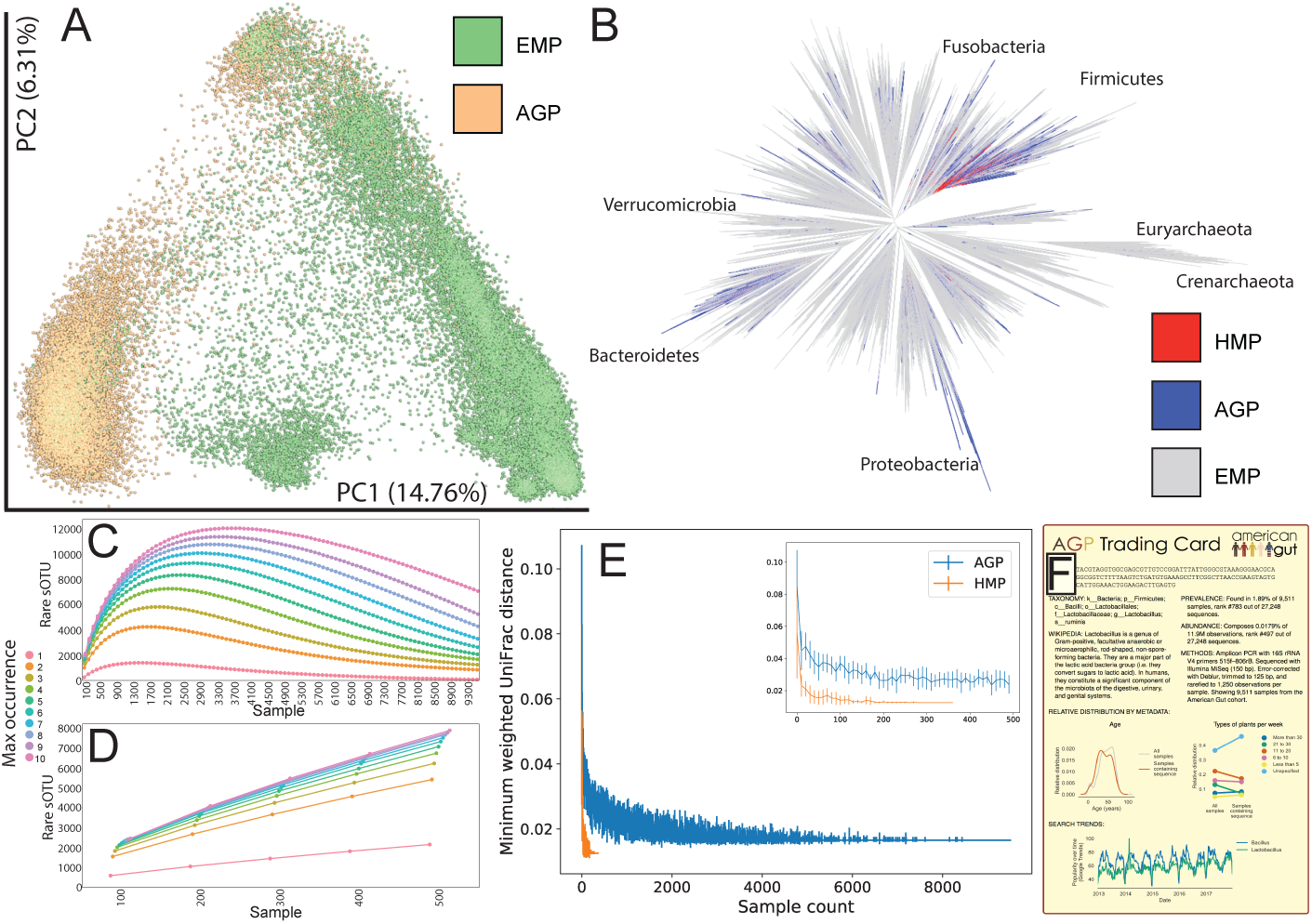
OTU and beta-diversity novelty. **(A)** The AGP data placed into the context of extant microbial diversity at a global scale. **(B)** A phylogenetic tree showing the diversity spanned by the AGP, and the HMP in the context of Greengenes and the EMP. **(C)** sOTU novelty over increasing numbers of samples in the AGP; the AGP appears to have begun to reach saturation and is contrasted with **(D)** Yatsunenko et al. 2012 which unlike the AGP had extremely deep sequencing per sample. **(E)** The minimum observed UniFrac distance between samples over increasing numbers of samples for the AGP and the HMP; inset is from 0-500 samples. **(F)** An AGP “trading card” of an sOTU of interest (shown in full in fig S2).

Inserting the sOTU fragments of AGP and EMP samples into a Greengenes (20) reference phylogenetic tree using SEPP (21) (fig 3B) showed that the AGP population harbored much broader microbial diversity than the Human Microbiome Project (5). Both datasets are dwarfed by the breadth of bacterial and archaeal phylogenetic diversity in environmental samples. Examining sOTUs over increasing numbers of samples, we observed a reduction in the discovery rate of novel sOTUs starting around 3,000 samples, emphasizing the need for focused sampling efforts outside the present AGP population (fig 3C). The importance of sample size for detecting novel microbes and microbiomes is apparent when contrasted against Yatsunenko et al. (6), which contained hundreds of samples from three distinct human populations at ∼1 million sequences/sample (fig 3D). This effect is magnified in beta-diversity analysis, where the AGP has saturated the configuration space, and new samples are not “distant” from existing samples (fig 3E). To encourage community engagement with sOTUs found in the AGP, we adapted the EMP “trading cards” for sOTUs (figs 3F, S2).

### Temporal and spatial analyses

Longitudinal samples are required for understanding human microbiome dynamics (22). We examined 565 individuals who contributed multiple samples and observed an increasing trend of intrapersonal divergence with time. Still, over time individuals resemble themselves more than others, even after one year (fig 4A).

**Figure 4.**
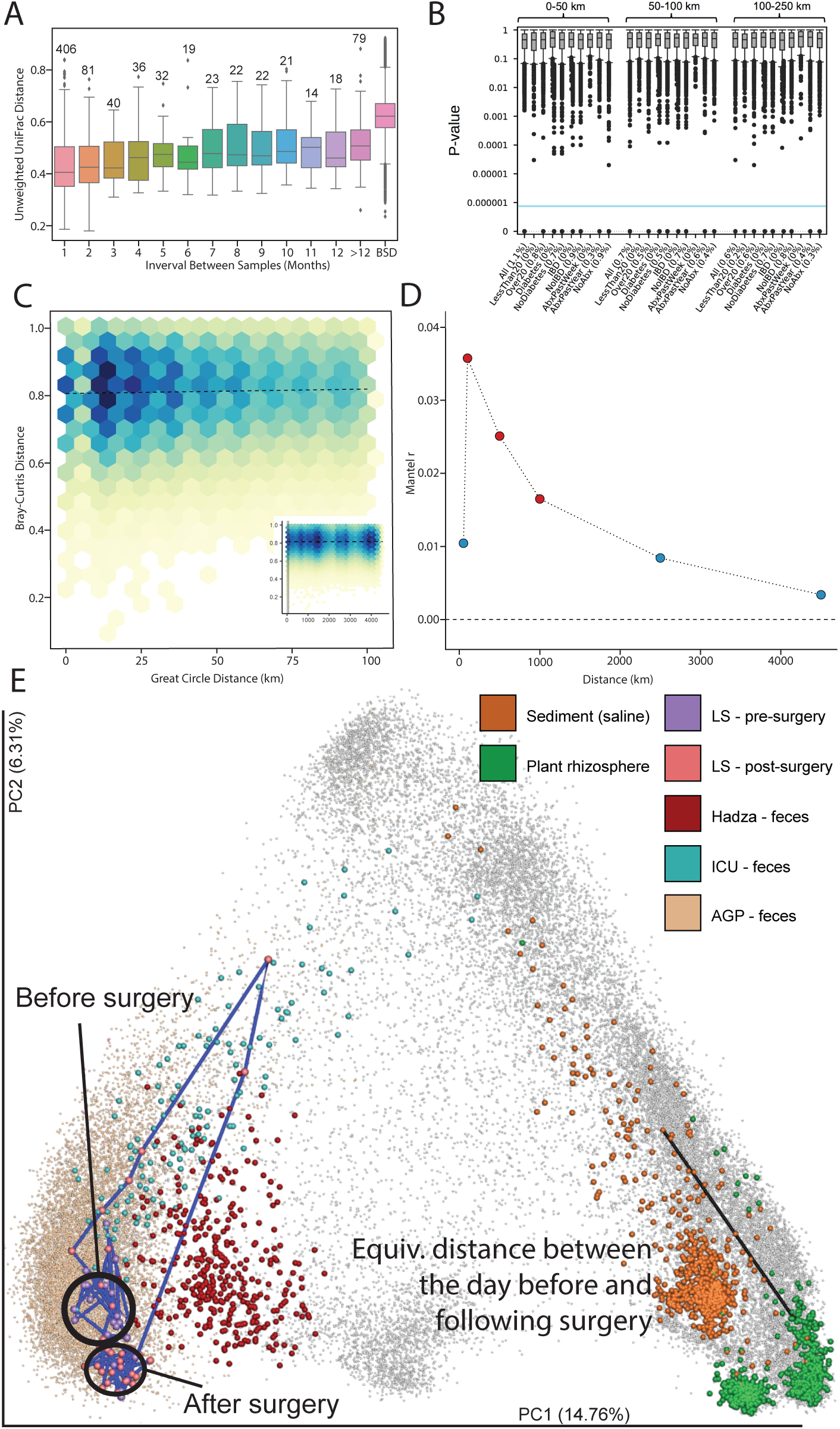
Temporal and spatial patterns. **(A)** 565 individuals had multiple samples. Distances between samples within an individual shown at 1 month, 2 months, etc out to over 1 year; between subject distances shown in “BSD.” Even at one year, the median distance between a participant’s samples is less than the median between participant distance. **(B)** Within the US, spatial processes of sOTUs appear driven by stochastic processes as few sOTUs exhibit spatial autocorrelation (Moran’s *I*) on the full dataset or partitions (e.g., participants older than 20). **(C)** Distance-decay relationship for Bray-Curtis dissimilarities between subject pairs that are within 100km (great-circle distance) radius of one another (Mantel test; *r*=0.036, adjusted *p*=0.03). Inset shows the largest radius (i.e., the contiguous US). Darker colors indicate higher-frequency bins. Dashed lines represent fits from linear models to raw data. **(D)** Mantel correlogram of estimated *r* coefficients, significance of distance-decay relationships, and radius (x-axis). Red points represent neighborhood sizes that were significant (adjusted *p*-values < 0.05). **(E)** Characterizing a large bowel resection using the AGP, the EMP, a hunter-gatherer population, and ICU patients in an unweighted UniFrac principal coordinates plot. A state change was observed in the resulting microbial community. The change in the microbial community immediately following surgery is the same as the distance between a marine sediment sample and a plant rhizosphere sample.

We tested whether patterns in individual longitudinal sample sets could be better explained when placed in the context of the AGP by integrating samples collected from: a) a time series of 58 time points from one subject (described as “LS”), prior to and following a large bowel resection, b) 2 time points from 121 patients in an intensive care unit (ICU) (23), c) samples from the “extreme” diet study from David et al. (24), and d) samples from the Hadza hunter-gatherers for additional context (25). Through the longitudinal sampling of LS, dramatic pre- and post-microbial configuration changes that exceeded the span of microbial diversity associated with the AGP population were observed (fig 4E, animated in (26)). After surgery, subject’s samples more closely resembled those of ICU patients (Kruskal Wallis H=79.774, p=4.197x^−19^, fig S2A-C), and showed a persistent state change upon return to the AGP fecal space. Remarkably, the UniFrac distance between the samples immediately prior to and following the surgery was almost identical to the distance between a marine sediment sample and a plant rhizosphere sample (unweighted UniFrac distance of 0.78). Furthermore, the observed state change in LS is not systematically observed in the extreme diet study (fig S2D; PERMANOVA n.s. when controlling for individual). Despite extensive dietary shifts, these subjects do not deviate from the background AGP context.

Recent reports suggest that the microbes of bodies (8), like those of homes (27), are influenced mostly by local phenomena rather than regional biogeography (28), and accordingly we observed only weak geographic associations with sOTUs (fig 4B), no significant distance-decay relationships (fig 4C), and, with Bray-Curtis distance, only a weak effect at neighborhood sizes of ca. 100km (Mantel *r*=0.036, Benjamini-Hochberg adjusted *p*=0.03) to 1,000km (Mantel *r*=0.016, Benjamini-Hochberg adjusted *p*=0.03).

### Dietary plant diversity

The self-reported dietary data suggested, unexpectedly, that the number of unique plant species a subject consumes is associated microbial diversity, rather than self-reported categories such as “vegan” or “omnivore” (fig 2D, E). Principal Components Analysis of FFQ responses (fig 5A) revealed clusters associated with diet types such as “vegan.” However, these dietary clusters did not significantly relate to microbiome configurations (fig 5B; Procrustes fig 5A, *M*^*2*^=0.988). We therefore characterized the impact of dietary plant diversity on the microbial community.

**Figure 5.**
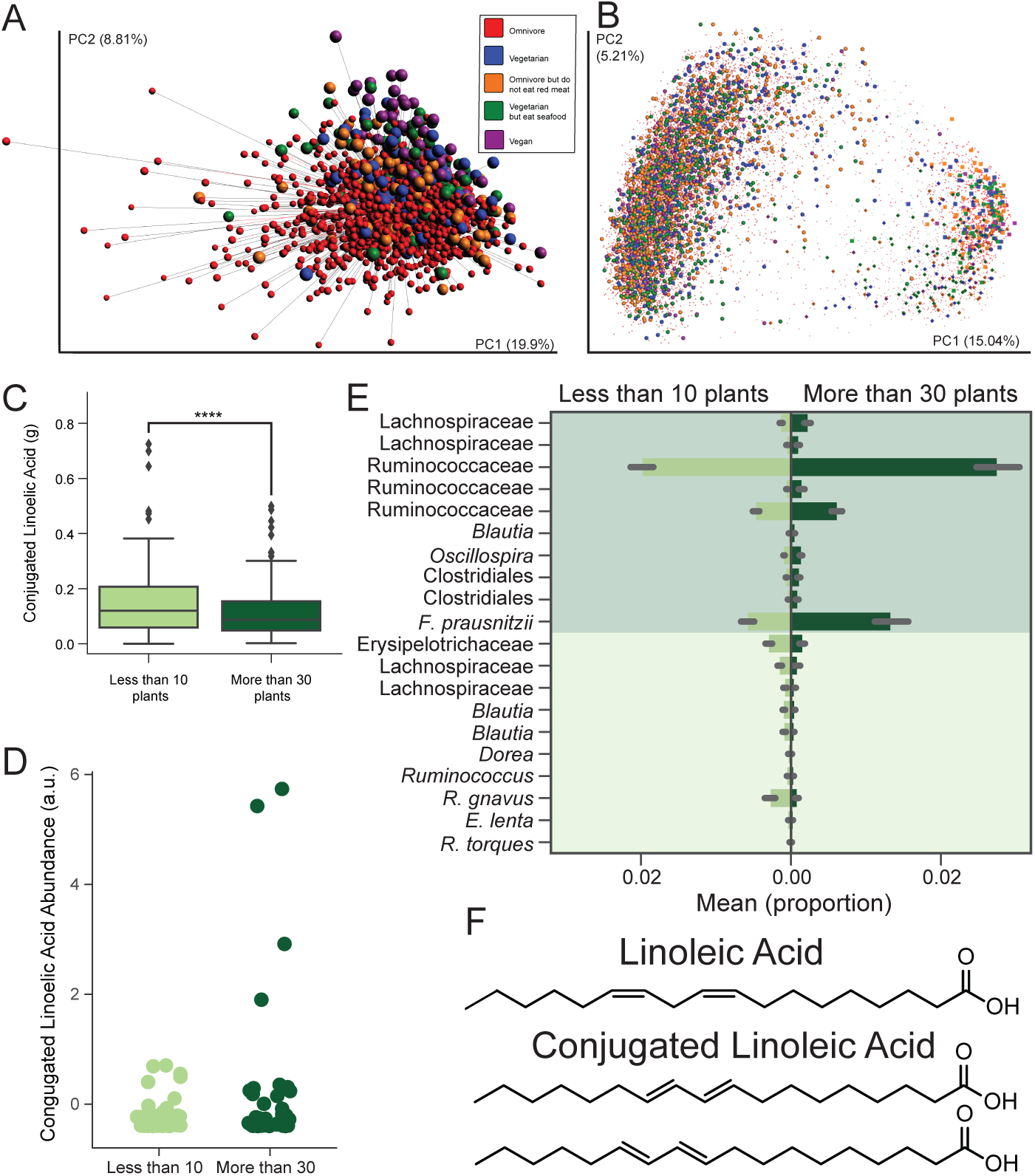
Diversity of plants in a diet. **(A)** Procrustes analysis of fecal samples from (*n*=1,596) individuals using Principal Components of the Vioscreen FFQ responses and Principal Coordinates of the unweighted UniFrac distances (*M*^*2*^=0.988) colored by diet; Procrustes tests the fit of one ordination space to another. PCA shows grouping by diets such as Vegan suggesting self-reported diet type is consistent with differences in micro and macro nutrients as recorded by the FFQ, however these dietary differences do not explain relationships between the samples in 16S space. **(B)** The full AGP dataset including skin and oral samples through unweighted UniFrac and Principal Coordinates Analysis highlighting a lack of apparent clustering by diet type. **(C)** Dietary conjugated linoleic acid levels as reported by the FFQ between the extremes of plant diversity consumption, and **(D)** the observed levels of CLA by HPLC-MS. **(E)** Differential abundances of sOTUs (showing the most specific taxon name per sOTU) between those who eat fewer than 10 plants per week vs. those who eat over 30 per week. **(F)** The molecules, linoleic acid (LA) and conjugated linoleic acid (CLA) (only trans-, trans-isomers are shown) were found to comprise the octadecadienoic acid found to be the key feature in this difference in number of plants consumption.

Using balances (10), we identified several putative short-chain fatty acid (SCFA) fermenters associated with eating more than 30 types of plants, including sOTUs putatively of the species *Faecalibacterium prausnitzii* and of the genus *Oscillospira* (29) (AUC=68.5%, ANOVA *p*=8.9×10^−39^, *F*=177.2) (fig 5E, table S3). These data suggest community-level changes associated with microbial fermentation of undigested plant components. Because bacteria differ in their carbohydrate-binding modules and enzymes that hydrolyze diverse substrates in the gut (30), a diet containing various types of dietary fibers and resistant starches likely supports a more diverse microbial community (31, 32).

To test these effects in the stool metabolome, we performed HPLC-MS annotation and annotation propagation (33, 34) on a subset of fecal samples (*n*=219) preferentially selecting individuals at the extremes of plant type consumption, i.e. eating <10 or >30 different types of plants per week. Several fecal metabolites differed between the two groups, with one key discriminating feature annotated as octadecadienoic acid (annotation level 2 according to the 2007 metabolomics initiative, (35)). Further investigation using authentic standards revealed that the detected feature was comprised of multiple isomers, including linoleic acid (LA) and conjugated linoleic acid (CLA). CLA abundance was significantly higher in individuals consuming > 30 types of plants, and those consuming more fruits and vegetables generally, (fig 5D, 1-sided *t*-test; p < 10^−5^), but did not correlate with dietary CLA consumption as determined by the FFQ (dietary fig 5C; Spearman *r* < 0.16; *p* > 0.15). CLA is a known end-product of LA conversion by lactic acid bacteria in the gut, such as *Lactobacillus plantarum* (36) and *Bifidobacterium* spp. (37). FFQ-based dietary levels of LA and MS-detected LA did not differ significantly between groups (fig S3), suggesting that their different microbiomes may differentially convert LA to CLA. Several other putative octadecadienoic acid isomers were also detected (fig 5F), some strongly correlated with plant consumption. Determining these compounds’ identities as well as their origin and function may uncover new links between the diet, microbiome, and health.

### Molecular novelty in the human gut metabolome

Our untargeted HPLC-MS approach allowed us to search for novel molecules in the human stool metabolome, parallel to our search for novelty in microbes and microbiome configurations described above. Bacterial N-acyl amides were recently shown to regulate host metabolism by interacting with G-protein-coupled receptors (GPCRs) in the murine gastrointestinal tract, mimicking host-derived signaling molecules (38). These agonistic molecules regulate metabolic hormones and glucose homeostasis as efficiently as host ligands. Manipulating microbial genes that encode metabolites eliciting host cellular responses could enable new drugs or treatment strategies for many major diseases, including diabetes, obesity, and Alzheimer’s disease: roughly 34% of all marketed drugs target GPCRs (39). We observed N-acyl amide molecules previously hypothesized but unproven to be present in the gut (38) (fig 6, S4), as well as new N-acyl amides (fig 6).

**Figure 6.**
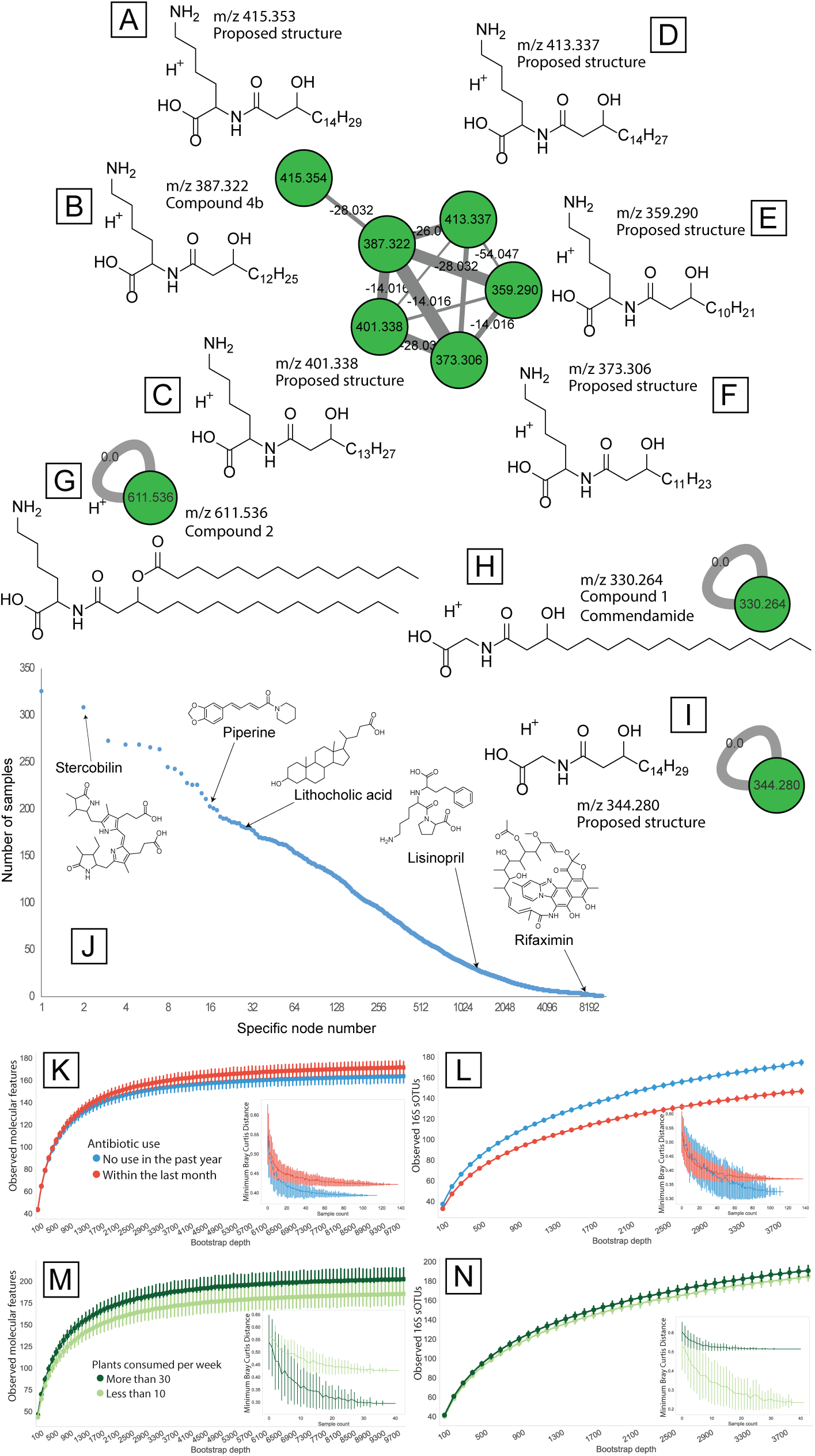
Molecular novelty in the gut microbiome. **(A-I)** Molecular sub-network of N-acyl amides. Cluster/nodes of microbially-derived G protein-coupled receptor agonistic molecules detected in human fecal samples are shown. Molecules B, G and H have been described (compounds 1, 2 & 4b (38) and commendamide (123)); molecules A, C, D, E and I are previously not reported (proposed structures are shown). **(J)** Compound occurrence frequency plot. Examples of compounds originating from food (piperine, black pepper alkaloid), host (stercobilin, heme catabolism product), bacterial activity (lithocholic acid, microbially-modified bile acid) or exogenous compounds such as antibiotics (rifaximin) or other drugs (lisinopril, high blood pressure medication) are shown. **(K-N)** Alpha and beta-diversity assessments of antibiotic and plants cohorts; insets depict minimum observed beta-diversity over increasing samples.

Levels of two N-acyl amides, annotated as commendamide (*m/z* 330.2635, fig S4B) and N-3-OH-palmitoyl ornithine (*m/z* 387.3220, fig S4C), positively correlated with a self-reported medical diagnosis of thyroid disease (Kruskal–Wallis, FDR *p*=0.032, *p*=2.48×10^−3^, *χ*2=11.99; N-3-OH-palmitoyl ornithine; Kruskal–Wallis, FDR *p*=0.048, *p*=5.63×10^−3^, *χ*2=10.35). Conversely, glycodeoxycholic acid (m/z 450.3187) was significantly higher in individuals not reporting thyroid disease diagnosis (Kruskal–Wallis; FDR *p*=1.28×10^−4^, *p*= 4.41×10^−7^, χ2=29.27). This cholic acid is produced through microbial dehydroxylation, again linking gut microbiota to endocrine function (40, 41).

Finally, we compared metabolome diversity to 16S diversity in the samples selected for dietary plant diversity and a second set of samples selected to explore antibiotic effects (*n*=256 individuals who self-reported not having taken antibiotics in the past year (*n*=117), or having taken antibiotics in the past month (*n=*139); participants were matched for age, BMI, and country). By computing a collector’s curve of observed molecular features in both cohorts (fig 6K, 6M), we observe that, paradoxically, individuals who had taken antibiotics in the past month (*n*=139) had significantly greater molecular diversity (Kruskal Wallis, *H*=255.240, *p*=1.87×10^−57^) than those who had not taken antibiotics in the past year (*n*=117), and differed in molecular beta-diversity (fig 6K inset), suggesting that antibiotics promote unique metabolomes that result from differing chemical and microbial environments in the gut. Notably, the diversity relationships of this set are not reflected in 16S diversity (fig 6L, 6N), where antibiotic use shows decreased diversity (Kruskal Wallis *H*=3983.839, *p*=0.0). Within the dietary plant diversity cohort, we observed a significant increase (Kruskal Wallis, *H*=897.106, *p*=4.17×10^−197^) in molecular alpha diversity associated with a high diversity of plant consumption (*n*=42) compared to low plant diversity (*n*=43), a relationship also observed in 16S diversity, where high dietary plant diversity increased 16S alpha diversity (Kruskal Wallis, *H*=65.817, *p*=4.947x^−16^).

### A living dataset

The AGP is dynamic, with samples arriving from around the world daily. This allows a living analysis, similar to continuous molecular identification and annotation revision in the Global Natural Products Molecular Networking (GNPS) database (34). Although the analysis presented here represents a single snapshot, samples continued to arrive during manuscript preparation. For example, after we defined the core “healthy” sample set, an exploratory analysis using matched controls was performed by collaborators to test for correlations between mental illness and microbiome composition (as reported in (42, 43)). By analyzing mental illness status (depression, schizophrenia, post-traumatic stress disorder (PTSD) and bipolar disorder – four of the most disabling illnesses per World Health Organization (44)) reported by AGP participants (*n*=125) against matched 1:1 healthy controls (*n*=125), we observed a significant partitioning using PERMANOVA in weighted UniFrac (*p*=0.05, *pseudo-F*=2.36). These findings were reproducible within US residents (*n*=122, p=0.05, *pseudo-F*=2.58), UK residents (*n*=112, *p*=0.05, *pseudo-F*=2.16), women (*n*=152, *p*=0.04, *pseudo-F*=2.35), and people 45 years of age or younger (*n*=122, *p*=0.05, *pseudo-F*=2.45). We also reproduce some previously reported differentially abundant taxa in Chinese populations using our UK subset (42, 45)(table S3). This shows that multi-cohort replication is possible within the AGP (additional detail supplemental text).

## Discussion

The AGP provides an example of a successful crowdfunded citizen science project that facilitates human microbiome hypothesis generation and testing on an unprecedented scale, provides a free data resource derived from over 10,000 human-associated microbial samples, and both recaptures known microbiome results and yields new ones. Ongoing living data efforts, such as the AGP, will allow researchers to document and potentially mitigate the effects of a slow but steady global homogenization driven by increased travel, lifespans, and access to similar diets and therapies, including antibiotics. Because the AGP is a subproject of the EMP (1), all samples were processed using the publicly available and widely used EMP protocols to facilitate meta-analyses, as highlighted above. Further example applications include assessing the stability of AGP runs over time, comparing the AGP population to fecal samples collected from a fecal transplant study (46) and an infant microbiome time series (47), the latter using different DNA sequencing technology, to highlight how this context can provide insight (48).

A unique aspect of the AGP is the open community process of assembling the Research Network and analyzing these data, which are released immediately on data generation. Analysis details are shared through a public forum (GitHub, https://github.com/knightlab-analyses/american-gut-analyses). Scientific contributions to the project were made through a geographically diverse Research Network represented herein as the American Gut Consortium, established prior to project launch and which has grown over time. This model allows a “living analysis” approach, embracing new researchers and analytical tools on an ongoing basis (e.g., Qiita (*Web:http://qiita.microbio.me*) and GNPS (34)). Examples of users of the AGP as a research platform include educators at several universities, UC San Diego Athletics, and the American Gastroenterological Association (AGA). Details on projects using the AGP infrastructure can be found in the supplement.

To promote public data engagement, we aimed to broaden the citizen science experience obtained by participating in AGP by “gamifying” the data and separately by developing an online forum for microbiome data discussion and discovery. The gamification introduces concepts of beta-diversity and challenges users to identify clusters of data in principal coordinates space (http://csb.cs.mcgill.ca/colonyb/). The forum, called Gut Instinct (http://gutinstinct.ucsd.edu), enables participants to share lifestyle-based insights with one another. Participants also have the option to share their AGP sample barcodes, which will help us uncover novel contextual knowledge. Gut Instinct now has over 1,050 participants who have collectively created over 250 questions. Participants will soon design and run their own investigations using controlled experiments to further understand their own lifestyle and the AGP data.

The AGP therefore represents a unique citizen-science dataset and resource, providing a rich characterization of microbiome and metabolome diversity at the population level. We believe the community process for involving participants from sample collection through data analysis and deposition will be adopted by many projects harnessing the power of citizen science to understand the world around and within our own bodies.

## Materials and methods

### Participant Recruitment and Sample Processing

Participants signed up for the project through Indiegogo (https://www.indiegogo.com/) and later, FundRazr (http://fundrazr.com/). A contribution to the project was made to help offset the cost of sample processing and sequencing (typically $99 per sample; no requirement to contribute if another party was covering the contribution). All participants were consented under an approved Institutional Review Board human research subjects protocol, either from the University of Colorado Boulder (protocol #12-0582; December 2012 - March 2015) or the University of California, San Diego (protocol #141853; February 2015 - present). The IRB-approved protocol specifically allows for public deposition of all data that is not personally identifying and for return of results to participants (fig. 1A).

Self-reported metadata were collected through a web portal (http://www.microbio.me/americangut). Samples were collected using BBL Culture Swabs (Becton, Dickinson and Company; Sparks, MD) and returned by mail. Samples were processed using the EMP protocols. Briefly, the V4 region of the 16S rRNA gene was amplified with barcoded primers and sequenced as previously described (49). Sequencing prior to August 2014 was done using the 515f/806r primer pair with the barcode on the reverse primer (50); subsequent rounds were sequenced with the updated 515f/806rB primer pair with the barcode on the forward read (51). Sequencing batches 1-19 and 23-49 were sequenced using an Illumina MiSeq; sequencing for 20 and 21 were performed with an Illumina HiSeq Rapid Run and round 22 was sequenced with an Illumina HiSeq High Output.

### 16S Data Processing

The 16S sequence data were processed using a sequence variant method, Deblur v1.0.2 (52) trimming to 125nt (otherwise default parameters), to maximize the specificity of 16S data; a trim of 125nt was used because one sequencing round in the American Gut used 125 cycles while the rest used 150. Following processing by Deblur, previously recognized bloom sequences were removed (14). The Deblur sub Operational Taxonomic Units (sOTUs) were inserted into the Greengenes 13_8 (53) 99% reference tree using SEPP (54). Taxonomy was assigned using an implementation of the RDP classifier (55) as implemented in QIIME2 (56). Multiple rarefactions were computed, with the minimum being 1250 sequences per sample with the analyses using the 1250 set except where noted explicitly. Diversity calculations were computed using scikit-bio 0.5.1 with the exception of UniFrac (57) which was computed using an unpublished algorithmic variant, Striped UniFrac (*https://github.com/biocore/unifrac)*, which scales to larger datasets and produces identical results to previously published UniFrac algorithms.

### Metadata Curation

To address the self-reported nature of the AGP data and ongoing nature of the project, basic filtering was performed on the age, height, weight, and body mass index (BMI). Height and weight were gated to only consider heights between 48 cm and 210 cm, and weight between 2.5 kg and 200 kg. BMI calculations using values outside this range were not considered. We assumed age was misreported by any individual who reported a birth date after their sample was collected. We also assumed age was misreported for participants who reported an age of less than 4 years, but height over 105 cm, weight over 20 kg, or any alcohol consumption. Values assumed to be incorrect were dropped from analyses (fig S1B).

### Sample Selection

Analyses in the manuscript were performed on a subset of the total AGP samples. A single fecal sample was selected for each participant with at least one fecal sample that amplified to 1250 sequences per sample unless otherwise noted. Priority was given to samples that were associated with VioScreen (*http://www.viocare.com/vioscreen.html*) metadata.

The samples used for analysis and subsets used in various analyses are described in table S2. Briefly, we defined the healthy subset (*n*=3,942) as adults aged 20-69 years with a BMI between 18.5 and 30 kg/m^2^ who reported no history of inflammatory bowel disease or diabetes and no antibiotic use in the last year. There were 1,762 participants who provided results for the VioScreen Food Frequency Questionnaire (FFQ; *http://www.viocare.com/vioscreen.html*). The meta-analysis with non-Western samples (*n*=4,643) included children over the age of 3, adults with a BMI of between 18.5 and 30 kg/m^2^, and no reported history of inflammatory bowel disease, diabetes, or antibiotic use in the last year.

### Population Level Comparisons

Population level comparisons were calculated for all American Gut participants living in the United States. BMI categorization was only considered for adults over the age of twenty, since the description of BMI in children is based on their age and sex. Education level was considered for adults over the age of 25. This threshold was used to match the available data from the US Census Bureau (*https://www.census.gov/content/dam/Census/library/publications/2016/demo/p20-578.pdf*). The percentage of the American Gut participants was calculated as the fraction of individuals who reported results for that variable. US population data is from the 2010 census (*https://www.census.gov/prod/cen2010/briefs/c2010br-03.pdf*), US Census bureau reports (*https://www.census.gov/content/dam/Census/library/publications/2016/demo/p20-578.pdf*), Centers for Disease Control reports on obesity (*https://www.cdc.gov/nchs/data/hus/2015/058.pdf*), diabetes (57, 58), IBD (*http://www.cdc.gov/ibd/ibd-epidemiology.htm*), smoking (*https://www.cdc.gov/tobacco/data_statistics/fact_sheets/adult_data/cig_smoking/index.htm*), and a report from the Williams Institute (*http://williamsinstitute.law.ucla.edu/wp-content/uploads/How-Many-Adults-Identify-as-Transgender-in-the-United-States.pdf*) (table S2).

### Within American Gut Alpha-and Beta-Diversity Analyses

OTU tables generated in the primary processing step were rarefied to 1,250 sequences per sample. Shannon, Observed OTU, and PD whole tree diversity metrics were calculated as the mean of ten rarefactions using QIIME (56, 59). Alpha-diversity for single metadata categories was compared with a Kruskal-Wallis test. Unweighted UniFrac distance between samples was tested with PERMANOVA (60) and permuted *t*-tests in QIIME.

### Balances

The goal of this analysis was to design two-way classifiers to classify samples and sOTUs. This will allow us to identify sOTUs that are strongly associated with a given environment. To do this while accounting for issues due to compositionality, we used balances (61) constructed from Partial Least Squares (62).

First the sOTU table was centered log-ratio (CLR) transformed with a pseudocount of 1. Partial least squares discriminant analysis (PLS-DA) was then performed on this sOTU table using a single PLS component, using a binary categorical variable as the response and the CLR transformed sOTU table as the predictor. This PLS component represented an axis, which assigns scores to each OTU according to how strongly associated they are to each class. An sOTU with a strong negative score indicates an association for the one category, which we will denote as the negative category. An sOTU with a strong positive score indicates that sOTU is strongly associated with the other category, which we will denote as the positive category.

We assumed that PLS scores associated with each OTU were normally distributed. Specifically

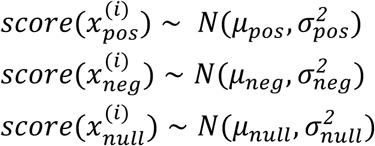

Where *μ*_*null*_ *≈ 0*, *μ*_*neg*_ < *0* and *μ*_*pos*_ > *0*. To obtain estimates of these normal distributions, Gaussian Mixture Models with three Gaussians were fitted from the PLS scores. Thresholds were determined from the intersection of Gaussians. The OTUs with PLS scores less than the intersection 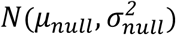 and 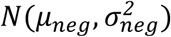 are classified to be associated with the negative category. The OTUs with PLS scores greater than the intersection 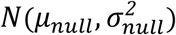 and 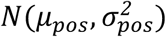 are classified to be associated with the positive category.

The balance was constructed as follows

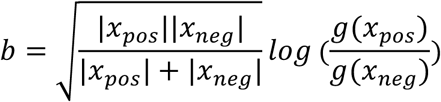

From this balance, we calculated receiver operator characteristic (ROC) curves and AUC to assess the classification accuracy, and ran ANOVA to assess the statistical significance. The dimensionality was shrunk through some initial filtering (an sOTU must have at least 50 reads, must exist in at least 20 samples except where noted, and have a variance over 10 to remove sOTUs that do not appear to change), so that the number of samples is greater than the number of sOTUs to reduce the likelihood of over-fitting. This technique was used to investigate differences due to plant consumption, country of residence and western vs non-western and was consistently applied with the exception that a filter of 5 samples was used for the western vs. non-western analysis due to group sample sizes.

Balances on plant consumption were constructed using Partial Least Squares. Only samples from people who consumed less than 10 types of plants a week or more than 30 types of plants a week were considered.

### Meta-analysis of samples from the American Gut and from individuals living agrarian and hunter-gatherer lifestyles

A meta-analysis compared fecal samples collected from healthy individuals that were 3 years of age or older and included in the AGP data set to a previously published 16S rRNA V4 region data set that included healthy people living an industrialized, remote agrarian or hunter-gatherer lifestyle (63-65). The AGP subset of healthy individuals was determined by filtering by the metadata columns “subset_antibiotic”, “subset_ibd”, “subset_diabetes”, and for individuals over the age of 16 years “subset_bmi”. All datasets were processed using the Deblur pipeline as noted above, with the exception that all reads in the meta-analysis, including AGP data, were trimmed to 100nt to accommodate the read length in Yatsunenko et al (63). Bloom reads as described above were removed from all samples. We used Striped UniFrac as noted above to estimate beta-diversity (unweighted UniFrac) and EMPeror software (66) version 0.9 to visualize principal coordinates. We used a non-parametric PERMANOVA with 999 permutations to test for significant differences in fecal microbiomes associated with industrialized, remote agrarian, and hunter-gatherer lifestyles. All AGP samples were considered to be from people living an industrialized lifestyle. Balances were constructed from Partial Least Squares to assess the differences between the hunter-gather vs. industrialized populations and the remote farmers vs industrialized populations.

### Spatial Autocorrelation

We sought to investigate distance-decay patterns – the relationship between microbial community similarity and spatial proximity – among American Gut participants, to determine the extent to which geographical distances could explain variation in microbial community taxonomic compositions between participant pairs. The correlation between community-level Bray-Curtis (67) distances and participants’ spatial proximities (i.e., great-circle distances, km) was assessed using a Mantel test (68) with 1000 matrix permutations. Analyses were conducted using the subset of participants located in the continental United States that had not received antibiotics in the last year. Different neighborhood sizes were investigated in order to detect the relevant spatial scale on which significant distance-decay patterns in microbial community compositions emerged. To accomplish this, we computed distance-decay relationships for a series of model adjacencies corresponding to neighborhood radiuses of 50, 100, 500, 1000, 2500, and 4500 km among participants, and adjusted *p*-values for multiple comparisons using the Benjamini-Hochberg procedure (69). We also studied spatial correlations in phylogenetic community dissimilarities, calculated as weighted normalized UniFrac distances, using the procedure described above. Analyses were conducted in R statistical programming environment.

The spatial autocorrelation of each individual taxon was assessed using Moran’s *I* statistic (70). Taxa present in less than 10 samples were filtered, since these would not be sufficiently powered. Analyses were conducted using binary spatial weight matrices, with neighborhoods of 0 – 50 km, 50 – 100 km, and 100 – 250 km. The different neighborhoods were useful for detecting spatial autocorrelation at different scales. All spatial weights matrices were row-standardized. We checked for spatial autocorrelation at three taxonomic ranks: class, genus, and OTU. We also considered whether there was autocorrelation within subsets of individuals who were under 20 years old and between 20 and 70 years old; those having IBD, no IBD, diabetes, and no diabetes; and those who had taken antibiotics within the past week, year, or not within the past year. The results presented above did not qualitatively depend on the subset of individuals considered. Statistical significance was assessed using permutation tests, which were implemented using a Markov Chain Monte Carlo algorithm. To assess each *p*-value, 100 chains were run each starting from a different random permutation. Each chain had 1000 iterations. We used Bonferroni corrections to correct for multiple comparisons, with an overall significance level set to 0.05. Analyses were run using custom Java code, optimized for running many spatial autocorrelation analyses on large data sets (71).

### Metadata cross-correlation

To account for covariance among metadata for effect size and variation analyses, we examined the correlation between individual metadata variables including technical parameters. Groups in ordinal variables were combined if there were insufficient sample size (e.g. people who reported sleeping less than 5 hours were combined with those who reported sleeping 5 to 6 hours into a variable described as “Less than 6”). The same transformations were used for effect size analysis. Any group with less than 25 total observations was ignored during analysis; if this resulted in a metadata column having no groups, the column was removed from analysis. The relationship between continuous and ordinal covariates was calculated using Pearson’s correlation. Ordinal and categorical covariates were compared using a modified Cramer’s V statistic (72). Continuous and categorical covariates were compared with a Welch’s T test (73). We treated used *1-R* as a distance between the covariates. Traversing the resulting binary, weighted cluster tree starting at tip level into the direction of the root, i.e. bottom-up, we grouped tips together that are members of the same subtree after covering a distance of approximately 0.5 (branch length 0.29). A representative variable from each cluster was selected for analysis (table S2).

### Effect Size Calculations

Effect size was calculated on 179 covariates (including technical parameters), selected from the cross-correlation (table S2). Ordinal groups with small sample sizes at the extreme were collapsed as noted above. Individuals who reported self-diagnosis or diagnosis from alternative practitioner for medical conditions were excluded from the analysis. Any metadata variable with less than 50 observations per group or that made up less than 3% of the total number of respondents was also excluded from the effect size analysis. Continuous covariates were categorized into quartiles. For each one of the 179 variables, we applied the mdFDR (74) methodology to test for the significance of each pairwise comparison among the groups. For each significant pairwise comparison, we computed the effect size using Cohen’s *d* (75), or the absolute difference between the mean of each group divided by the pooled standard deviation. For analysis of diversity, we used Faith’s Phylogenetic Diversity (alpha-diversity) and weighted and unweighted UniFrac distances (beta-diversity).

### Variation analysis

Using the methodology reported in the supplemental material for (76), we computed Adonis (77) using 1000 permutations, over the sample sets used in the effect size calculations as noted above, and applied Benjamini-Hochberg correction (FDR<0.1) to assess drivers of variation in beta-diversity.

### Meta-analysis movie

American Gut samples from all body sites were combined with data from an infant time series (78), a fecal transplant study (79), and recent work characterizing the microbiome of patients in the intensive care unit (80). The combination of the datasets in movie S2 required that all sequences were trimmed to an even length of 125 nucleotides. All projects except for the infant time series were sequenced using an Illumina instrument. In order to combine the data, we expressed the Illumina and non-Illumina data through a common reference database. Specifically, the Deblur sOTUs from the Illumina data were mapped against the Greengenes (53) database (13_8 release) using 99% similarity; the associations between the input sOTUs, and their cluster memberships, were used to construct an OTU table based on the original sOTU per sample sequences counts (i.e., summing the counts for all sOTUs in a common OTU). The infant time series data were picked using a closed reference OTU picking approach against the same reference at the same similarity. The infant time series dataset followed a closed reference OTU picking approach using 99% similarity. The resulting two tables (from Illumina-generated data and the ITS dataset) were merged and analyzed using the Greengenes 99% tree. The table was rarefied to 1,250 sequences per sample. Principal coordinates projections were calculated based on unweighted UniFrac distance (57). The principal coordinates analysis was visualized and animated in EMPeror 1.0.0-beta8-dev (66, 81). The movie was captured in QuickTime (Apple, Cupertino, CA), and edited with Premiere Pro (Adobe, San Jose, CA).

### Integration with the Earth Microbiome Project

A precomputed 100nt Deblur BIOM table representing the data in (82) was obtained from (ftp://ftp.microbio.me/emp/release1/otu_tables/deblur/). 100nt Deblur tables were also obtained from Qiita for Hadza fecal samples (Qiita study ID 11358, (83)), ICU microbiome samples (Qiita study ID 2136, (80)), and a longitudinal series which includes samples immediately prior to and following a large bowel resection (Qiita study ID 10283, EBI accession ERP105968, unpublished); all samples were processed using the EMP Illumina 16S V4 protocol. The EMP dataset used a minimum sOTU count of 25; the same threshold was applied to the other datasets included prior to merge. Blooms as identified by (84) were removed from all samples. This collection of BIOM tables was then merged yielding an OTU table representing 40,600 samples. sOTUs were restricted to those already present in the EMP 100nt fragment insertion tree, which represents 329,712 sOTUs. The table was then rarefied to 1000 sequences per sample, and unweighted UniFrac computed using 768 processors with the aforementioned Striped algorithm. Visualizations and animations were performed using EMPeror v1.0.0b12.dev0.

### Extreme diet study state assessment

The sequence data from (85) were processed by Deblur to assess 16S sOTUs in common with the AGP processing above. In order to assess a state difference with PERMANOVA, we needed to control for sample independence within the longitudinal sampling. To do so, we randomly selected one sample from each individual per diet, computed PERMANOVA, and repeated the process 100 times. None of the trials produced a *p*-value below 0.05.

### Vioscreen PCA and diet type Procrustes analysis

Before performing Principal Component Analysis (PCA) on the informal diet questions, Vioscreen variables that are categorical or receive less than 90% response among the 1762 participants were excluded leaving 1596 participants. PCA was then performed using the Vioscreen information from these participants’ responses over 207 Vioscreen questions, and then colored by their types of diet as answered in the AGP informal food survey. The coordinates from the PCA were extracted. For the same samples, PCoA of unweighted UniFrac distances was computed on the 16S data subset from the primary processing set. The coordinates from the PCA and the PCoA were assayed for a measure of fitness using Procrustes as implemented in QIIME v1.9.1.

### Beta-diversity added

To assess added beta-diversity, we applied the technique used in (86) figure 3. Specifically, we randomly sampled *N* samples from the distance matrix 10 times, over an increasing value of *N*. For each set of sampled distances, we computed the minimum observed distance.

### sOTU novelty

To assess sOTU novelty, we randomly sampled *N* samples from an sOTU table 10 times, over an increasing value of *N*. At each sampling, we computed the number of sOTUs observed with read counts within minimum thresholds. In other words, a minimum threshold of 1 is the number of singletons observed in the sampled set, a minimum threshold of 2 is the number of singletons and doubletons, etc.

### Within-individual beta-diversity

Many of the individuals in the American Gut Project contributed multiple samples, but at uneven time intervals. In order to explore intrapersonal variation, we replicated the analysis in Lloyd-Price et al. figure 3 (87). Specifically, we determined all time deltas between a subjects samples, and gathered the distributions of beta-diversity between any two samples binned by month. An individual is only represented a single time in a given month, but may be represented in multiple months if they had, for instance, contributed samples over the course of a year.

### High Performance Liquid Chromatography Mass Spectrometry (HPLC-MS) Analysis

A total of 498 samples were selected for analysis via mass spectrometry. Specifically, two groups were chosen. First, given the large body of primary literature describing the negative impact of antibiotics on the gut microbiome, and the general interest in this topic from many American Gut participants, we chose 279 samples from individuals (age, BMI, and country matched) who self-reported not having taken antibiotics in the past year, or having taken antibiotics in the past month or week. We chose a second group of 219 samples collected from individuals who answered the question “In an average week, how many different plants do you eat? (e.g., if you consume a can of soup that contains carrots, potatoes and onion, you can count this as 3 different plants; If you consume multi-grain bread, each different grain counts as a plant. Include all fruits in the total)” on the main American Gut Project main survey and who had also completed the VioScreen Food Frequency Questionnaire. When American Gut participants collect samples, they do so on a double headed swab; therefore, all samples chosen for this analysis had one remaining swab head (the first had been used for DNA extraction and microbiome sequencing).

#### Cell cultures sample preparation for metabolomics analysis

The supernatant collected from cell cultures (see “expanded bloom assessment” below) were processed to make them compatible with HPLC-MS analysis. The solid phase extraction with wash was carried out to reduce impact of cell culture media, which is highly detrimental for the ESI. The 30 mg sorbent Oasis HLB (Waters, Waltham, MA) SPE cartridges were used to achieve broad metabolite coverage. The cell samples were stored at −80°C and thawed at room temperature immediately prior to extraction. The thawed samples were then centrifuged for 10 minutes at 1200 rpm and extracted. For the SPE extraction, the Oasis HLB SPE cartridge was conditioned with 700µL of 100% HPLC-grade methanol and equilibrated with 700µL of HPLC-grade DI water. The cell supernatant (∼350-400µL) was loaded into cartridge and allowed to slowly elute. The loaded SPE wells were then washed with 800µL of 5% methanol in water and the absorbed material was slowly eluted with 200 µL of 100% methanol. Vacuum up to ∼ 20 psi was applied for the wells that did not elute within an hour. The collected eluent was stored at −20°C until the HPLC-MS analysis.

#### Fecal sample preparation for metabolomics analysis

The swab tubes scheduled for analysis were removed from the −80°C freezer and placed on dry ice for the duration of sample processing. Each tube with swab was logged by reading the barcode with barcode scanned and the swab was removed from tube and placed onto a ThermoFisher Scientific (ThermoFisher Scientific, Waltham, MA) 2 ml deep well 96-well plate set on top of dry ice coolant. The top part of each swab’s stick was snapped off and discarded. Immediately after filling all of the wells with swabs, 200 µL of HPLC-grade 90% v:v ethanol:water solvent was added to each well using multichannel pipette. Four blanks of unused swabs and extraction solvent were included onto each plate. Each plate was then sealed with 96-well plate lid, sonicated for 10 minutes and placed into the refrigerator at 2 °C to extract samples overnight. After extraction, the swabs were removed from wells and discarded, the plates were placed into a lyophilizer, and the entire sample was dried down and then re-suspended in 200 µL 90% v:v ethanol:water. The plates were resealed and centrifuged at 2000 rpm for 10 minutes. The 100 µL aliquots of sample were then transferred onto a Falcon 96-well MS plate using a multichannel pipette, and each plate was immediately sealed with sealing film. The MS plates were centrifuged at 2000 rpm for 10 minutes and stored at 2 °C until analysis.

#### HPLC-MS analysis

The metabolomics analysis of samples was conducted using reverse phase (RP) high performance liquid chromatography mass spectrometry (HPLC-MS). The HPLC-MS analysis was performed on a Dionex UltiMate 3000 ThermoFisher Scientific high-performance liquid chromatography system (ThermoFisher Scientific, Waltham, MA) coupled to a Bruker impact HD qTOF mass spectrometer. The chromatographic separation was carried out on a Kinetex C18 1.7 µm, 100Å UHPLC column (50 mm x 2.1 mm) (Phenomenex, Torrance, CA), held at 40 °C during analysis. A total of 5 µL of each sample was injected. Mobile phase A was water, mobile phase B was acetonitrile, both with added 0.1% v:v formic acid. The solvent gradient table was set as follows: initial mobile phase composition was 5% B for 1 min, increased to 40% B over 1 min, then to 100% B over 6 min, held at 100% B for 1 min, decreased back to 5% B in 0.1 min, followed by a washout cycle and equilibration for a total analysis time of 13 min. The scanned m/z range was 80-2000 Th, the capillary voltage was 4500 V, the nebulizer gas pressure was 2 bar, the drying gas flow rate was 9 L/min, and the temperature was 200 °C. Each full MS scan was followed by MS/MS using collision-induced dissociation (CID) fragmentation of the seven most abundant ions in the spectrum. For MS/MS, the collision cell collision energy was set at 3 eV and the collision energy was stepped 50%, 75%, 150% and 200% to obtain optimal fragmentation for differentially sized ions of different sizes. The scan rate was 3 Hz. A HP-921 lock mass compound was infused during the analysis to carry out post-processing mass correction. All of the raw data are publicly available at the UCSD Center for Computational Mass Spectrometry (*111*) (dataset ID: MassIVE MSV000080179).

#### MS data analysis

The collected HPLC-MS raw data files were first converted from Bruker’s *d* to mzXML format and then processed with the open source OpenMS 2.0 software (88) in order to deconvolve and align each peak across different chromatograms (feature detection). The alignment window was set at 0.5 minutes, the noise threshold at 1000 counts, the chromatographic peak FWHM value at 20, and the mass error at 30 ppm. All of the peaks that were present in any of the blanks with S/N below 10:1 were removed from the final feature table. The number of features with corresponding MS/MS was as follows: Vioscreen study sample cohort: 5144 total MS2 features; antibiotics study samples cohort: 8288 total MS2 features. The number of MS1 features is difficult to estimate exactly as it depends on feature detection settings and the number of samples, but it is typically about 4-5 fold greater depending on the sample. For all of the MS1 features detected across all samples, only ∼1-5% are present in an individual sample.

Chemical annotations were carried out by automatic matching fragmentation spectra to multiple databases using Global Natural Product Social Molecular Networking (GNPS) (89) and then examining the data at the MS/MS level by molecular networking (90). The goal is to retrieve spectra with identical and similar fragmentation patterns and combine them into consensus nodes and clusters, respectively. The consensus node spectra are then compared against public MS/MS libraries to provide molecular annotations (91). Further annotations could be suggested by examining the molecular network (90) (so called propagated annotations). Annotations obtained with precursor and MS/MS matching are considered level two annotations according to the 2007 metabolomics standards initiative (92). All molecular networking analysis and annotations are available here: antibiotic use subset (93); types of plants subset (94), cell cultures of isolates (95) and fecal samples co-networked with the cell cultures (96). The raw data contain a significant number of abundant features originating from swab polymers. Therefore, selective background peak removal was carried out specifically for the polymer compounds originating from swabs that were used for the sample collection. The m/z shifts that correspond to the polymer repeating units (44.0262, 88.0524, 132.0786, 176.1049) were identified with GNPS m/z differences frequency plot. The network clusters that contained nodes with the corresponding mass differences were deemed to belong to polymers and all member nodes of the network clusters were removed from the feature table (a total of 1632 features/nodes). Principal Coordinates Analysis (PCoA) using a Hellinger distance (97) matrix was used to confirm that the batch effect corresponding to the batches of swabs was mitigated prior to further analysis. To confirm putative annotations, authentic standards were purchased for the linoleic acid (LA; Spectrum Laboratory Products, Inc., USA), conjugated linoleic acid (CLA; mixture of 4 isomers: 9,11 and 10,12 isomers, E and Z) (Sigma-Aldrich, USA), and selected antibiotics: tetracycline, oxytetracycline, and doxycyclin (Abcam Inc., USA). For level one identifications, each authentic compound was analyzed under identical experimental conditions and retention time and MS/MS spectra were compared with putatively annotated compounds.

#### Selective feature detection

Selective feature extraction was performed with open source MZmine2 software (98). To separate closely eluting LA and CLA isomers as well as separate various N-acyl amides, crop filtering with RT range of 5.4-6.0 minutes and m/z range of 281.246 - 281.248 was applied to all chromatograms. Mass detection was performed with a signal threshold of 1.0E2 and a 0.6 s minimum peak width. The mass tolerance was set to 20 ppm and the maximum allowed retention time deviation was set to 5 s. For chromatographic deconvolution, the baseline cutoff algorithm with a 5.0E1 signal threshold was used. The maximum peak width was set to 0.5 min. Similarly, the MS feature for reference compound stercobilin was extracted with a crop filter RT range of 2.0-4.0 minutes and m/z range of 595.345-595.355. The stercobilin reference compound was used to assess variability of chromatographic retention times to ensure that the compounds of interest (LA and CLA in particular) retention times were correctly identified. After isotope peak removal, the peak lists of all samples were aligned within the corresponding retention time and mass tolerances. Gap filling was performed on the aligned peak list using the peak finder module with 1% intensity, 10 ppm m/z tolerance, and 0.05 min RT tolerance, respectively. After the creation and export of a feature matrix containing the feature retention times, exact mass, and peak areas of the corresponding extracted ion chromatograms, we added sample metadata to the feature matrix metadata of the samples.

The selective feature extraction with the same settings has been performed for all of the detected compounds listed on the Figure 6A-I (the m/z range crop filter window was set for corresponding m/z for each compound).

#### Molecular Networking

Raw data files were converted to the .mzXML format using Bruker Data Analysis software and uploaded to the GNPS (https://gnps.ucsd.edu/) MassIVE mass spectrometry database (https://massive.ucsd.edu/). Molecular networking was performed to identify spectra shared between different sample types and to identify known molecules in the data set. All annotations are at level 2 according to the proposed minimum standards in metabolomics (92). The data were filtered by removing all MS/MS peaks within +/-17 Da of the precursor m/z. MS/MS spectra were window-filtered by choosing only the top 6 peaks in the +/-50 Da window throughout the spectrum. The MS spectra were then clustered with MS-Cluster algorithm with a parent mass tolerance of 0.02 Da and a MS/MS fragment ion tolerance of 0.02 Da to create consensus spectra (89). Further, consensus spectra that contained less than 4 spectra were discarded. A network was then created where edges were filtered to have a cosine score above 0.65 and more than 5 matched peaks. The edges between two nodes were kept in the network if and only if each of the nodes appeared in each other’s respective top 10 most similar nodes. The spectra in the network were then searched against GNPS spectral libraries. The library spectra were filtered in the same manner as the input data. All library matches were required to have a score above 0.7 and at least 6 matched peaks. Molecular networks were visualized and mined using the Cytoscape software (www.cytoscape.org/).

#### Molecular networking-based propagation of annotations

The annotation of GPCR agonist compounds was not possible via direct library matching, as their spectra are not present in any MS libraries, but direct comparison with fragmentation patterns presented in (99) allowed us to establish these compounds’ identity with level 3 identification (92). Consequently, manual annotation of compounds was carried out in two steps. The exact mass of compounds and their MS/MS fragmentation spectra were matched to the reference spectra found in supplementary info of (99) (fig S4A). Compound m/z 611.5357 was identified in this fashion. In addition, commendamide (330.2640) and its analogue (m/z 344.2799) were identified by matching exact mass of the corresponding ion and by in silico prediction of the MS/MS fragmentation spectra with the CSI:FingerID (100) (fig S4B). For novel molecules that were found within clusters of compounds of interest, but were not described in the literature previously, the structure was postulated using annotation propagation from adjacent annotated nodes in the cluster as described in (89) by assessing differences in parent mass and fragmentation patterns. The key structure, m/z 387.322 has been annotated as N-3-OH-palmitoyl ornithine based on the exact mass and previous annotation (99) as well as analysis of fragmentation pattern to confirm structural moieties of fragments (fig S4C). The rest of the structural assignments have been propagated from that structure. The ornithine moiety has been determined to be present in each structure (due to presence of the signature ion with m/z 115.09), and acylation of the hydroxyl is not possible due to insufficient mass of the structures; thus, the changing mass was postulated to correspond to different length of the alkyl substituent (fig 6, in the main text).

#### Correlations of Metabolites with Metadata

We have investigated correlations between metabolites (especially those of interest, such as N-Acyl amides) and all of the categories in the metadata. The data were subsetted into the Vioscreen and Antibiotics cohorts and normalized using probabilistic quotient normalization (101). In order to test the association of the metabolites to the categorical metadata fields we performed the Kruskal–Wallis test followed by Benjamini & Hochberg FDR correction to all metabolites. The significant metabolite-metadata associations (*p*-value adjusted < 0.05) were further connected to GNPS spectral library matches associating the MS1 feature to the MS2 precursor ion in a 10 ppm mass window and 20 seconds retention time window. The results are summarized in table S5.

#### Data pretreatment for statistical analysis

A PCoA plot using Hellinger distance (distance matrix: Hellinger; grouping: HCA) was built with all samples in the subset; one sample was found to be an outlier and removed. The data were then filtered to remove features with near-constant, very small values and values with low repeatability using the inter-quartile range estimate. Detailed description of methodology is given in (102). The samples were normalized by sum total of peak intensities, an important step due to large variability of the fecal material load on different swabs. To reduce the effect of background signal and make the sum normalization appropriate, the subtraction of blank and polymer peak features was conducted prior to analysis, as described above. The data were further scaled by mean centering and dividing by standard deviation for each feature.

The data were split into two groups for downstream analysis. Group one contained samples from individuals answering “More than 30” (*n*=41) and “Less that 10” (*n*=44) to the main American Gut Project survey question “In an average week, how many different plants do you eat?” Group two contained samples from individuals answering “antibiotic use within last week” (*n*=56) and “I have not taken antibiotics in the past year” (*n*=115) to the main American Gut Project survey question “I have taken antibiotics in the last —.” for the Antibiotic history study, correspondingly.

The resultant features tables were used as input for the Metabonalist software (103). Partial least squares Discriminant Analysis (PLS-DA) (62) was used to explore and visualize variance within data and differences among experimental categories. Random forests (104) (RF) supervised analysis was used to further verify validity of determined discriminating features.

### Expanded bloom assessment

The American Gut Project dataset now spans multiple-omics types, and include data that were unavailable during the analysis described in Amir et al. (14). To better understand how the blooming organisms impacted the samples in the American Gut, we 1) performed an additional set of 16S-based experiments; 2) cultured historical samples covering a range of bloom fractions, characterized their metabolites and sequenced the isolates; 3) performed shotgun metagenomics sequencing on the “high bloom” samples; 4) ran the set of samples previously run for HPLC-MS (e.g., the plants and antibiotics cohorts) for shotgun metagenomics, and 5) ran the storage samples from (105) for shotgun metagenomics. The additional sequencing effort was to provide a basis to assess whether functional potential driven by the blooms was impacting any of the biological results discussed in the manuscript. The additional HPLC-MS work was to characterize the metabolites specific to the blooms to remove them from analysis. The additional sequence data generated from the American Gut samples were deposited in EBI under the American Gut accession (ERP012803), and the storage sample data under its accession (ERP015155).

#### 16S-based bloom experiments

Effect size calculations were computed prior to and following the removal of bloom reads using the procedure described by Amir et al., 2017 (84). The fraction of reads recruiting to blooms was included as a covariate. Effect sizes were assessed over Faith’s Phylogenetic Diversity (59), unweighted UniFrac (57) and weighted UniFrac (106). We then computed Pearson and Spearman correlations of the effect sizes, per metric, between the bloom and bloom-removed result (fig 2D, E). In addition to the effect size calculations, we also tested whether the bloom fraction was correlated to any metadata category and did not observe significant correlations.

We then tested the removal of blooms from other studies in which room temperature shipping was not performed by retrieving a wide variety of human fecal studies from Qiita. UniFrac distance matrices were computed prior to and following bloom removal, followed by Mantel tests. The results of this procedure are outlined in table S4.

Finally, we correlated the relative intensities of the HPLC-MS data associated with the antibiotics and plants cohorts against the fraction of blooming reads. Critically, we observed a set of spectra that are significantly correlated (table S5) to this fraction. On annotation using molecular networking (discussed in detail the HPLC-MS section), we observed these metabolites to putatively be LysoPE, lysophospholipid (LPL), which has previously been associated with the release of colicin (107). These metabolites were removed from subsequent analyses.

#### Culturing

Primary specimens (*n*=214) were selected from three plates based off of the median fraction of reads recruiting to the blooms across the plate, whether the primary specimen still existed, and as to gather samples from at least the US (*n*=116) and UK (*n*=73); additional countries were included in smaller sample sizes and include Australia (*n*=7), Germany (*n*=7), Canada (*n*=3), Croatia (*n*=2), Belgium (*n*=2), France (*n*=1), Austria (*n*=1), Sweden (*n*=1), and the Czech Republic (*n*=1). The bloom typically observed in these samples (and in the full AGP dataset) is an *E. coli* (ID: 04195686f2b70585790ec75320de0d6f from (84)), although a few of the other bloom sequences were represented at high read fraction as well. Samples were retrieved from −80°C and thawed on ice. The swab head was broken off into 500 µl sterile 1x Dulbecco’s Phosphate-Buffered Saline and vortexed vigorously for 30 seconds. Serial dilutions from this initial stock were made including 1:10,000 and 1:1,000,000. 10µl of the 1:10,000 dilution were inoculated into 1.5 ml sterile Tryptic Soy broth (TSB, BD cat#2253534) in sterile 96-deep-well plates (community cultures, CC) and incubated overnight at 37°C on an orbital shaker at 500 rpm. OD600 values above 0.1 (TSB controls measured ∼0.08) were counted as positive growth. Samples with high bloom fraction tended to grow overnight in ambient conditions, samples with a low bloom fraction tended to not grow in these conditions (fig 2A). Additionally, 100 µl of each dilution were plated onto Tryptic Soy agar using sterile glass beads and incubated overnight at 37°C. The following morning, a picture of the best dilution was captured and the most representative colony was selected from each plate and inoculated into 1.5 ml sterile TSB for overnight incubation as above (isolates, IS). The following morning, OD600 measurements were taken and the cultures were pelleted at 3,000 g for 5 min. The supernatant and cell pellets were stored at −20°C for metabolomic analysis and DNA extraction, respectively.

Shotgun sequencing was performed on all isolates and community cultures using a 1:10 miniaturized Nextera library prep with 1 ng gDNA input or up to 1 µl and a 15 cycle PCR amplification. Libraries were quantified with PicoGreen^TM^ dsDNA Assay Kit and 50 ng of each library (or 4 µl maximum) was pooled. The library was size-selected for 200-700 bp using the Sage Bioscience Pippin Prep and sequenced as a paired end 150 cycle run on an Illumina HiSeq 2500 v2 in Rapid Run mode at the UCSD IGM Genomics Center. Sequence processing including assembly performed as in the metagenomic processing section below with the exception that “-- meta” was not used with SPAdes (108), and read binning against the resulting contigs was not performed. For each isolate, contigs with abnormally high or low coverage as defined by the 1.5 × IQR rule were dropped. The characterization of the metabolites from the supernatant using HPLC-MS is discussed in the HPLC-MS section above.

Following assembly of the draft genomes, taxonomic assessment by Kraken (109) revealed that of the 119 successfully sequenced colony isolate cultures, 95 matched the bloom organisms identified by Amir et al., 2017. Compellingly, 70 of these isolate genomes contained exact 16S sequence matches to a bloom organism identified by (84), including 65 of which matched the dominant *E. coli* bloom in the American Gut (table S4).

The read data for the isolates were then assessed for predicted biosynthetic gene clusters (BGCs). We used biosyntheticSPAdes (110) to analyze BGCs in the assembly graph of individual genomes. Below we focus on the longest BGCs that are particularly difficult to reconstruct based on ad hoc analysis of contigs and reveal their variations (that likely translate into variations of their natural products). Some of the reconstructed long BGC are ubiquitous (shared by many isolates, albeit with some variations), while others are unique, e.g., present in a single or small number of isolates. We identified BGCs, representing in the alphabet of their domains (table S4), and uncovered variations in their sequence across multiple isolates. Specifically, a ubiquitous BGC similar to the elusive peptide-polyketide genotoxin colibactin and a unique surfactin-like BGC. Colibactin triggers DNA double-strand breaks in eukaryotic cells (111, 112) and induces cellular senescence and metabolic reprogramming in affected mammalian cells (113). Of the 11 samples containing the longest colibactin-like BGC, 10 of them contained the exact *E. coli* bloom 16S sequence described above; the 11^th^ isolate was actually a canine fecal sample plated alongside human (as the AGP allows participants to submit pet samples).

Although colibactin is frequently harbored by various *E. coli* strains, the variations of colibactin BGCs across various isolates have not been studied before. Genomic analysis revealed wide variations in colibactin-like BGCs suggesting that various strains produce related but not identical variants of natural products (114). These variations may give rise to the suite of LysoPE-associated spectra identified between the 16S and HPLC-MS datasets.

#### Shotgun sequencing of the high bloom and storage samples

Previously extracted DNA from the “high bloom” samples used for culturing was obtained, as was previously extracted DNA from Song et al. (105). Shotgun sequencing libraries from a total of 5 ng (or 3.5 µl maximum) gDNA was used in a 1:10 miniaturized KAPA HyperPlus protocol with a 15 cycle PCR amplification. Libraries were quantified with PicoGreen^™^ dsDNA Assay Kit and 50 ng (or 1 µl maximum) of each library was pooled. The pool was size-selected for 300-700 bp and sequenced as a paired end 150 cycle run on an Illumina HiSeq 2500 v2 in Rapid Run mode at the UCSD IGM Genomics Center. Sequence processing including assembly was performed as in the metagenomic processing section below.

#### Functional assessment of conjugated and non-conjugated linoleic acid

To investigate the metabolic potential of gut microbiome for producing conjugated linoleic acid from linoleic acid, we estimated the abundance of linoleic acid isomerase (LAI) in the fecal metagenome. We focused this investigation on the “plants” cohort, which were samples selected to maximize the difference between the number of types of plants metadata category as discussed in the main text. First, we translated the assembled metagenomes to metaproteomes using Prodigal gene prediction software. To map LAI to these metaproteomes, we used a representative LAI protein sequence (UniProt: D2BQ64), which was matched against UniProtKB (via https://www.ebi.ac.uk/Tools/hmmer/) for multiple sequence alignment (MSA). The resulting MSA file in clustal format was then used to generate a hidden Markov model (HMM) profile for LAI using hmmbuild in HMMER software (115). Subsequently, we mapped the resulting HMM profile to sample metaproteomes using hmmsearch with an E-value threshold of 10E-5. We calculated abundances of LAI per sample based on abundance (coverage x length) of LAI containing contigs in each sample, normalized to total sample biomass and performed linear regression between LAI abundances and bloom fraction. We did not note any correlation between metabolic potential of gut metagenome to produce LAI and the fraction of blooming bacteria (samples with no LAI hits were removed from this analysis). Similarly, there was no correlation between CLA abundances and bloom fraction in the samples. These results suggest that our report on the differential abundance of CLA in subjects with different dietary practices (with respect to the number of different types of plants consumed) is unlikely to be confounded by the presence of blooming bacteria.

#### Storage sample assessment

Metagenomic reads from the storage samples were mapped to the 169 isolate assemblies. We then ran model comparison tests on each to determine which mappings were significantly different between frozen samples and samples left out at ambient temperatures for various periods of time. Using the ‘lme’ package (116) in R (v3.3.3. R Core Team 2017), linear mixed effects models were applied to the abundances, with individual treated as the random effect. Mappings were considered to be significantly associated with temperature if the model was significantly improved (ANOVA *p*<=0.05) by incorporating a fixed effect of temperature. Seven mappings to isolates were found to be significantly increased in samples stored in ambient temperatures compared to frozen samples in both storage studies, of which 3 contained the 16S of the dominant *E. coli* bloom in the AGP samples, and 2 contained the 16S from other blooms recognized by (84).

#### Shotgun sequence processing

Raw FastQ files were processed using Atropos v1.1.5 (117) to remove adapters and low-quality regions. Putative human genome contaminations were identified and removed by using Bowtie2 v2.3.0 (118) with the “--very-sensitive” option against the human reference genome GRCh37/hg19.

Sequences were assigned taxonomy using Kraken v1.0.0 (109) against the “standard” database built following the Kraken manual, which contains all complete bacterial, archeal, and viral genomes available from NCBI RefSeq as of Aug. 3, 2017. Results were processed using Bracken v1.0.0 (119) to estimate the relative abundance of species-level taxa.

Metagenome sequencing data were assembled using SPAdes v3.11.1 (108) with the “-- `meta`” flag enabled. Contigs ≥ 1 kb in length were retained and fed to the prokaryotic genome annotation pipeline Prokka v1.12 (120). putatively individual genomes were inferred using MaxBin2 v2.2.4 (121).

In parallel, contigs were sheared into 200-bp fragments and taxonomy was assigned using Kraken (see above). For each contig, the most assigned taxon at each taxonomic rank and the proportion of sequences assigned to it was inferred.

A total of 3725 genome bins were identified from 677 out of 780 AGP metagenomes, with 5.50 ± 4.05 bins per sample, and a maximum bin number of 30. Bins with completeness < 50% were dropped, leaving 1029 bins from 464 samples (2.22 ± 1.97 bins per sample, maximum bins = 19).

### Filtering Bacterial Blooms for Metabolomics Analysis

To assess and account for the impact of the metabolites contributed by these organisms, we have performed HPLC-MS analysis of cultures of blooming organisms to establish possible contributions, as described above. The raw data are publicly available at the UCSD Center for Computational Mass Spectrometry (*http://massive.ucsd.edu/*, dataset ID: MassIVE MSV000081777). It was found that there is a negligible overlap of the bloom-associated metabolites with the compounds detected in AGP samples (fig 2B). Furthermore, we have verified that none of the compounds discussed in this work (LA, CLA, compounds on Fig 6A-I) are present in these bloom cultures. The main organism implicated in bloom was determined to be *E. coli*, as described earlier and MS data corroborate these findings (fig 2C).

Considering that the metabolites resulting from microbial activity in cultures can differ significantly from those in vivo (e.g. many of the metabolites could originate not from de novo synthesis, but rather from microbial modifications of external compounds that are not present in media, e.g. from the host), we also explored associations of metabolites in AGP metabolomics samples and blooms. Spearman rank correlation analysis of the fraction of 16S reads in a sample reporting as bloom to metabolites observed in the same samples revealed several features that correlate significantly (table S5). There exists a significant overlap between the Antibiotics and Vioscreen studies subsets, indicating potential common origin of these features. The strongest correlation was found for the feature m/z 480.3106 with multiple bloom organisms (*ρ^∧^*2 > 0.25 for *E. coli* at *p* < 1e-40). This feature was found to also significantly correlate with the principle coordinates of the PCoA, with and without blooms in the UniFrac matrices for both subsets. The tentative annotation of this feature is lysoPE, a lysophospholipid (LPL). The LPLs production in vivo is a result of phospholipase A enzymatic activity associated with Gram-negative bacteria. It is known that lysoPE is essential for release of colicin (107). Colicin (by itself not detectable with the MS methodology in this study due to very high molecular mass) is a bacteriocin related to microbial warfare and is known to be produced by *E. coli*, the major bloomer in AGP. It can be suggested that the blooming of an organism is related to attempting to kill competitors to maximize nutrient availability. Importantly, removal of all of the features associated with bloom does not alter the metabolomics results at all, which indicates that all of the observed biological trends reported here are not related to blooms.

### Mental health in the American Gut Project

From AGP cohort, we selected subjects who endorsed a mental health disorder (depression, schizophrenia, PTSD, and/or bipolar disorder). This resulted in 1,140 subjects. 636 subjects endorsed at least one of the exclusion criteria (antibiotic use in the last year, IBD, *C. difficile* infection, pregnancy, Alzheimer’s, anorexia or bulimia, history of substance use disorder, epilepsy or seizure disorder, kidney disease, phenylketonuria). Out of the remaining 504 subjects, 319 did not provide information regarding country of residence, hence forming a case cohort of 185 subjects. The remaining samples were further filtered down to 125 samples to include only high quality fecal microbiome data (at least 1,250 sequences/sample) at a single time point per subject. For those cases, we created a 1:1 matched sample of patients and non-psychiatric comparison (NC) participants based on age (±5 years), BMI, history of diabetes, smoking frequency, country of residence, census region (if in US), and sequencing plate. For each of the cohorts we calculated beta-diversity distance matrices using Bray-Curtis dissimilarity and weighted UniFrac. On resulting matrices we ran pairwise PERMANOVA with 999 permutations between “cases” (people who reported mental illness) and NCs (out matched control dataset). Differential abundance testing was performed using permutive mean difference test at 10,000 permutations, with discrete FDR (122) correction at alpha=0.1.

## Acknowledgments

The authors would like to thank first all the citizen science volunteers around the world. We also thank Katherine Amato, Margaret K. Butler, Gabriela Surdulescu, Luke Ursell, Jeff DeReus, Evguenia Kopylova, John Graham, Victoria Vazquez, Jai Ram Rideout, Merete Eggesbo, Steve Green, Kathy Holt, Jim Huntley, Zehra Esra Ilhan, Daniel Mayer, Catherine Nicholas, Laura Parfrey, Juanma Peralta, Dorota Porazinska, Jennifer Smilowitz, Matt Gebert, Se Jin Song, Kumar Thurimella, Sam Way, Tony Walters, Sophie Weiss, Ulla Westermann, Shawnelle White, Doug Woodhams, Jerry Kennedy, Scott Handley, Chandni Desai, Carolina Carpenter, Gabriella Fanelli, Deborah L. Bright and Xochitl C. Morgan for their contributions to the American Gut Project. Some sequencing was conducted by Mahdieh Khosroheidari and Clifford L. Green at the IGM Genomics Center, University of California, San Diego, La Jolla, CA.The de-identified participant and sequence data for the American Gut are available from the European Bioinformatics Institute under accession number ERP012803; the HPLC-MS data are available in MassIVE under accession MSV000080179. Per sample accession numbers are included in the supplemental information (table S1). The EBI accession includes more samples than presented here as some samples were consented after the analysis was completed. Crowdfunding support was provided through Indiegogo and FundRazr. Some consortium members were supported by NSF IGERT award 1144807, the Sloan Foundation Microbiology of the Built Environment Program, the Wyss Institute, the Howard Hughes Medical Institute, the U.S. Dept. of Energy under Contract DE-AC02-06CH11357.NS, the ERC Starting Grant 2013 (European Research Council, Starting grant 336452-ENIGMO), WELBIO WELBIO-CR-2012S-02R and Robert Tournut award (Société Nationale Française de Gastro-Entérologie). The British Gut collection was carried out by the Department of Twin Research at KCL funded by the Wellcome Trust, MRC, JPI Dinamic grant and the National Institute for Health Research (NIHR) Clinical Research Facility at Guy’s & St Thomas’ NHS Foundation Trust and NIHR Biomedical Research Centre based at Guy’s and St Thomas’ NHS Foundation Trust and King’s College London. Tim Spector is an NIHR senior Investigator. This research was performed in accordance with the University of Colorado Boulder’s Institutional Review Board protocol #12-0582 and the University of California San Diego’s Human Research Protection Program protocol #141853. Parts of the computations were performed using the San Diego Supercomputer Center (SDSC) through XSEDE allocations, which is supported by the NSF grant ACI-1053575. We thank the BioFrontiers Institute for their administrative support for the project while it was based at the University of Colorado Boulder.

## Supplementary Text

**Effect size comparisons**

**Multi-cohort replication detail**

**Projects using the American Gut infrastructure**

**American Gut Survey**

**Supplemental references**

## Supplementary Figures

**Figure S1.** Workflow and population scale analyses. **(A)** Heatmap of income levels from the US Census and American Gut participant locations. **(B)** Sample flowchart for what sample sets correspond to each analysis. **(C)** Using PLS-DA we observed separation between US (*n*=6,634) and UK (*n*=2,071) fecal samples. **(D)** We performed a Principal Coordinates analysis comparing children over the age of 3 and adults from industrialized (*n*=4,643 AGP samples, *n*=4,927 samples total), remote farming (*n*=131), and hunter-gatherer (*n*=30) lifestyles.

**Figure S2.** Trading cards and LS’s samples compared to ICU patients and AGP participants and diet state change analysis. **(A)** Unweighted UniFrac distance distributions for the sample immediate prior to surgery vs. all ICU fecal samples, and distances of the sample immediately following surgery vs. all ICU fecal samples (Kruskal Wallis *H*=79.774, *p*=4.198x^−19^). **(B)** Same as panel **(A)** except comparing against all AGP fecal samples (Kruskal Wallis *H*=8117.734, *p*=0.0). **(C)** The median distances of each sample in Larry’s longitudinal dataset compared to both ICU and AGP. The last pre-surgery sample is on day 25 and the first post-surgery sample is day 27. **(D)** A principal coordinates analysis of UniFrac distances of the American Gut Project, samples from the “extreme” diet study by David et al. (85), and the Earth Microbiome Project. No obvious state change by the diet of the participants in David et al. is observed.

**Figure S3.** Dietary levels of linoleic acid based on validated food frequency questionnaire responses, and the detected linoleic acid by mass spectrometry did not differ significantly between groups consuming few or many types of plants per week.

**Figure S4**. Metabolomic identification and annotation. **(A)** Manual annotation via comparison of experimental MS fragmentation patterns to those given in (99). Top panel: reference spectrum for the “Compound 2” in (99); bottom panel: experimental MS/MS spectrum for the parent ion m/z 611.5357. The compound is annotated as 3-(myristoyloxy)palmitoyl lysine. **(B)** I*n silico* annotation using CSI:FingerID (100) for the ion with m/z 330.2640. Top panel: experimental fragmentation pattern explained by the putative fragmentation tree; bottom panel: the possible candidate structures ranked by match %. The top structure with 71.02% match corresponds to commendamide. **(C)** Manual annotation via comparison of experimental exact mass to that of identified compound in (100), N-3-OH-palmitoyl ornithine. The peaks in experimental MS/MS spectrum are examined and compared to theoretical fragments that would result from breaking bonds in the proposed structure. The structure is deemed to be consistent with the N-3-OH-palmitoyl ornithine annotation.

## Supplemental Tables

**Table S1.** Summary of sample numbers and type in the American Gut other studies, sample distributions by country and territory, sample distributions by US state, US participant demographics and per sequencing round sample accessions in EBI.

**Table S2.** American Gut data dictionary, proportion of responses per AG survey question that are represented as a single question; multiselect responses were omitted as these are stored in the metadata as per response type, informal dietary questions and correlations to the food frequency questionnaire, effect size results without bloom sOTUs, variable mapping with Falony et al. 2016 Science.

**Table S3.** sOTUs relevant to the balance analyses, and summary of differentially abundant taxa in UK cohort (negative effect size indicated the taxon is more prevalent in control (NC) subjects).

**Table S4.** Application of the filter for blooms to other human fecal studies which were not subjected to room temperature shipping, taxonomy of the draft isolate genomes, the specific bloom 16S sOTUs observed, and ubiquitous colibactin-like biosynthetic gene clusters (top) and a unique surfactin-like biosynthetic gene cluster observed in the bloom isolates.

**Table S5.** A set of molecular features which appeared to significantly correlate to the bloom fraction, and Kruskal–Wallis tests for metabolites in the Antibiotics and Vioscreen cohorts of samples.

